# Os-circANK negatively regulates resistance to bacterial blight in rice via Osa-miR398b/OsCSD1/OsCSD2 pathways

**DOI:** 10.1101/2024.06.18.599603

**Authors:** Xiaohui Liu, Peihong Wang, Sai Wang, Weixue Liao, Mingyan Ouyang, Sisi Lin, Rongpeng Lin, Panagiotis F. Sarris, Vasiliki Michalopoulou, Xurui Feng, Zinan Zhang, Zhengyin Xu, Gongyou Chen, Bo Zhu

**Author notes:** Correspondence: Bo Zhu.

## Abstract

CircRNAs are widely present in plants, yet there have been no reports on the role of circRNA in the regulatory mechanisms of interactions between rice and pathogenic bacteria. As the sequence of circRNAs matches the parent gene except for the back splicing site, there is no feasible way to specifically knockdown circRNAs in plants. We identified a circRNA, named Os-circANK, derived from an ankyrin repeat protein, which was significantly downregulated post *Xanthomonas oryzae* pv. *oryzae* (*Xoo*) inoculation. We have created a CRISPR-Cas13d system to specifically knockdown a plant circular RNA, resulting in genetically edited rice with a targeted decrease in Os-circANK expression. Os-circANK was found to function as a sponge for Osa-miR398b, suppressing the cleavage of *OsCSD1/OsCSD2* by Osa-miR398b, leading to a reduction in ROS levels following *Xoo* infection and a negative regulation of resistance to bacterial blight in rice. Our findings demonstrate that Os-circANK inhibits the resistance to bacterial blight in rice via the Osa-miR398b/OsCSD1/OsCSD2 pathways.

## Introduction

Circular RNAs (circRNAs), a recent addition to the non-coding RNA family, emerge as novel entities characterized by a covalently closed circular structure formed through reverse splicing events, distinguishing them from the majority of linear RNAs (Huang et al., 2023). Research indicates that circRNAs exhibit structural stability, diverse categories, sequence conservation, and specific expression patterns in cells and tissues (Hansen et al., 2013). They harbor numerous potential functions and play pivotal roles in governing gene expression and various biological processes (Liu and Chen, 2022). With the advancement of high-throughput sequencing technology and bioinformatics, thousands of circRNAs have been identified in various plants (Hong et al., 2020). The expression of plant circRNAs also responses to both biotic and abiotic stresses, including pathogen invasion (Chen et al., 2018; Gao et al., 2019; Chen et al., 2021). In plants, prior research has primarily concentrated on the identification and annotation of potential circRNAs, along with investigations into their functions. Nevertheless, the functional linkage between circRNA production and expression changes and plant disease resistance remains unknown.

The gain or loss of function in transgenic plants is essential for studying the functionality of circRNAs. The “reverse complementary assisted exonic splicing strategy” has been employed to achieve the overexpression of plant circRNAs (Gao et al., 2019) and has been applied in numerous studies involving plant growth and development, as well as responses to biotic and abiotic stress (Gao et al., 2023). The overexpression of CircR5g05160 in rice has been shown to enhance resistance against the rice blast fungus (Fan et al., 2020). The overexpression of grape Vv-circATS1 has been reported to enhance cold tolerance in Arabidopsis (Gao et al., 2019). However, distinguishing the function of circRNA from its homologous linear mRNA is challenging due to the shared overlapping sequences, aside from the back-splicing junction (BSJ) (Kristensen et al., 2019; Han et al., 2020). Therefore, reducing the levels of circRNA without affecting the expression of its parent gene remains a challenge (Kristensen et al., 2019). For instance, Zhou et al. employed the CRISPR-Cas9 system to knockout the rice Os03circ00204, and observed a significant decrease in the expression of its parent gene in transgenic plants (Zhou et al., 2021). Consequently, there is currently no feasible method for the specific knockdown of plant circRNAs.

Several studies in animals have demonstrated that circRNAs can sequester miRNAs, leading to the upregulation of their target genes (Hansen et al., 2013; Guo et al., 2014). The initial report on plant circRNAs by Wang et al. in 2014 (Wang et al., 2014) sparked interest in unraveling the connection between circRNAs and miRNAs in plants. Moreover, there are reports confirming the interaction between plant miRNAs and circRNAs. Following the overexpression of miR160a in poplar protoplasts (Liu et al., 2019) or miR156 in *Phyllostachys edulis* protoplasts (Wang et al., 2019a), the expression levels of circRNAs with predicted targeting relationships exhibited a significant decrease. Bioinformatics analysis predicted numerous putative OsMIR408 binding sites for Os06circ02797, and downregulation of seven out of nine hypothesized OsMIR408 target genes was observed in the os06circ02797Δ1 mutant, indicating that Os06circ02797 might function as a sponge for OsMIR408 (Zhou et al., 2021). In grapes, Vv-circSIZ1 has been observed to alleviate the repressive effect of VvmiR3631 on its target *VvVHAc1* (Gao et al., 2023). However, direct evidence and systematic studies proving the mutual interaction between circRNA and miRNA have not been reported to date.

Bacterial Leaf Blight (BLB) is one of the most prevalent bacterial diseases affecting rice in the southern regions of China (Nino-Liu et al., 2006). It is caused by the pathogenic strain *Xanthomonas oryzae* pv. *oryzae* (*Xoo*). Severe outbreaks of BLB can lead to a rice yield reduction of up to 50% under field conditions, posing a serious threat to rice food security (Nino-Liu et al., 2006). Current research on BLB and other common diseases in rice primarily focuses on coding genes (Busungu et al., 2016), and there is a relative lack of studies on the functional role of circRNAs in the interaction between rice and pathogenic bacteria. In this study, we focused on a differentially expressed circular RNA (Wang et al., 2022), Os-circANK, post Xoo infection, to investigate its functional characteristics in the rice-*Xoo* interaction. Overexpression and knockdown experiments of Os-circANK revealed its negative regulation on rice resistance to *Xoo*. Furthermore, Os-circANK was found to enhance the expression of Osa-miR398b target genes *OsCSD1* and *OsCSD2* by sponging Osa-miR398b, subsequently reducing reactive oxygen species (ROS) levels, thereby negatively modulating rice resistance to *Xoo*.

## Results

### Identification and Verification of Os-circANK

Based on the previous research (Wang et al., 2022), we found a OSA_circ12938 that is formed by the back splicing of exon2 of an ankyrin repeat-containing protein and contains 917 nucleotides (Fig. 1a), which was identified and named Os-circANK in this study. To determine the junction sequence of Os-circANK, PCR analysis was conducted using both genomic DNA (gDNA) and cDNA as templates, with convergent and divergent primers. Notably, PCR products were observed for both gDNA and cDNA templates when using convergent primers. However, PCR product was only generated when using cDNA template and divergent primers (Fig. 1a). The back-spliced junction was successfully amplified and verified by Sanger sequencing (Fig. 1a). The RNase R digestion assay confirmed that the linear form of the OsANK gene was sensitive to RNase R, whereas Os-circANK was resistant due to its loop structure (Fig. 1b). To validate the findings of previous studies on the differential expression of Os-circANK post *Xoo* infection^20^, we examined Os-circANK expression in Nipponbare plants. Our results showed a gradual decrease in Os-circANK transcription at 2, 6, and 12 hours post-inoculation, followed by a slight recovery at 24 hours. Nevertheless, the overall expression levels of Os-circANK were significantly decreased post *Xoo* inoculation (Fig. 1c). Both of the nucleoplasmic fractionation assay and FISH experiment consistently demonstrated that Os-circANK was abundantly expressed in the cytosol (Fig. 1d, e), which suggests that Os-circANK could function as molecular sponge of miRNA or RBP.

**Fig. 1.**
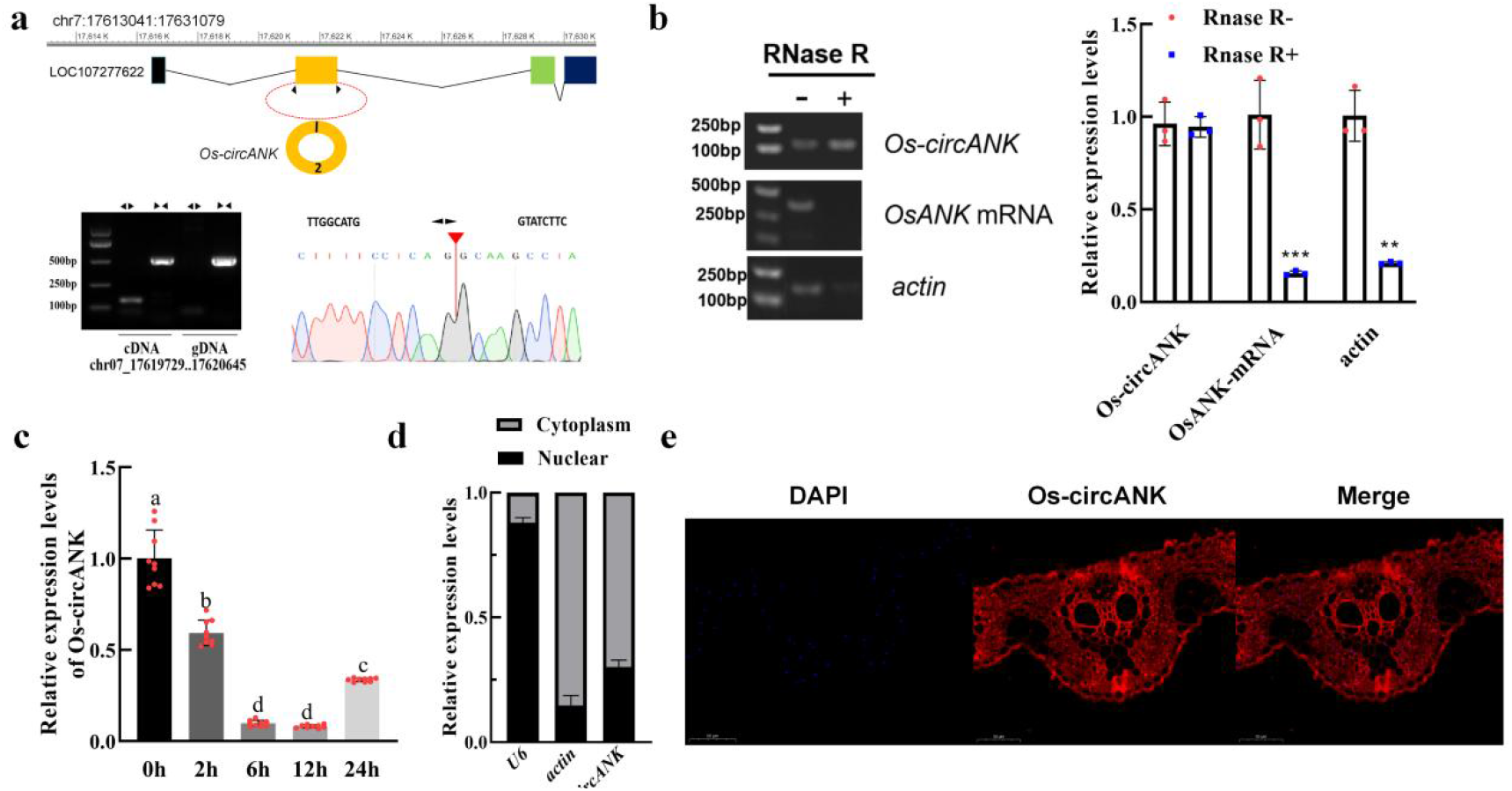
Characteristics of Os-circANK. a. Validation of Os-circANK by PCR using divergent primers. Upper panel, a diagram showing Os-circANK structure and the positions of back splicing site. Lower left panel, a pair of divergent primers (◀ ▶ ) amplified the Os-circANK within the cDNA, but not within the genomic DNA (gDNA); a pair of convergent primers (▶ ◀ ) was used as a control. Lower right panel, an example of Sanger sequencing illustrates the exon-derived back splicing of Os-circANK. b. The stability of Os-circANK was assessed through RNase R resistance assay and RT-qPCR analysis of both Os-circANK and OsANK, with and without RNase R treatment. *actin* served as a reference gene, mean±SD, t-test, \*\**p* < 0.01, \*\*\**p* < 0.001. c. RT-qPCR analysis of Os-circANK in leaves of the Nipponbare rice upon *Xoo* treatment. Different letters above the bars indicate significant differences at *p* < 0.01. d. Detection of Os-circANK in the cytoplasmic and nuclear fractions by RT-PCR (n = 3). e. Determination of the localization of Os-circANK by FISH using confocal microscopy. Os-circANK was stained in red, and nuclei were stained with DAPI (blue). All of the experiments were repeated three times with similar results.

### Efficient knockdown of rice circRNA with CRISPR/Cas13d system

We designed and optimized a plant-codon-optimized RfxCas13d protein, incorporating it into the pCAMBIA1300 vector backbone. Subsequently, we successfully synthesized the AtU6-crRNA gene fragment, which encompassed two distinct *Bsa* I enzyme cleavage sites, and subsequently established the CRISPR/RfxCas13d system, as illustrated in Fig. 2a. Finally, we devised gRNA targeting sequences spanning the BSJ region, each with a length of 15 nucleotides (Fig. 2b). Via agrobacterium-mediated transformation, rice plants with targeted downregulation of Os-circANK were acquired. The qPCR analysis revealed a notable reduction in the expression of Os-circANK compared to the control, whereas the mRNA expression of its corresponding parental genes exhibited an increase (Fig. 2c). These observations signify the specificity of the CRISPR/RfxCas13d vector system in selectively downregulating the expression of rice circRNAs. The increased expression of the parental gene following the knockdown of the circANK suggests that circANK may modulate the expression of its parental gene through a regulatory mechanism.

**Fig.2.**
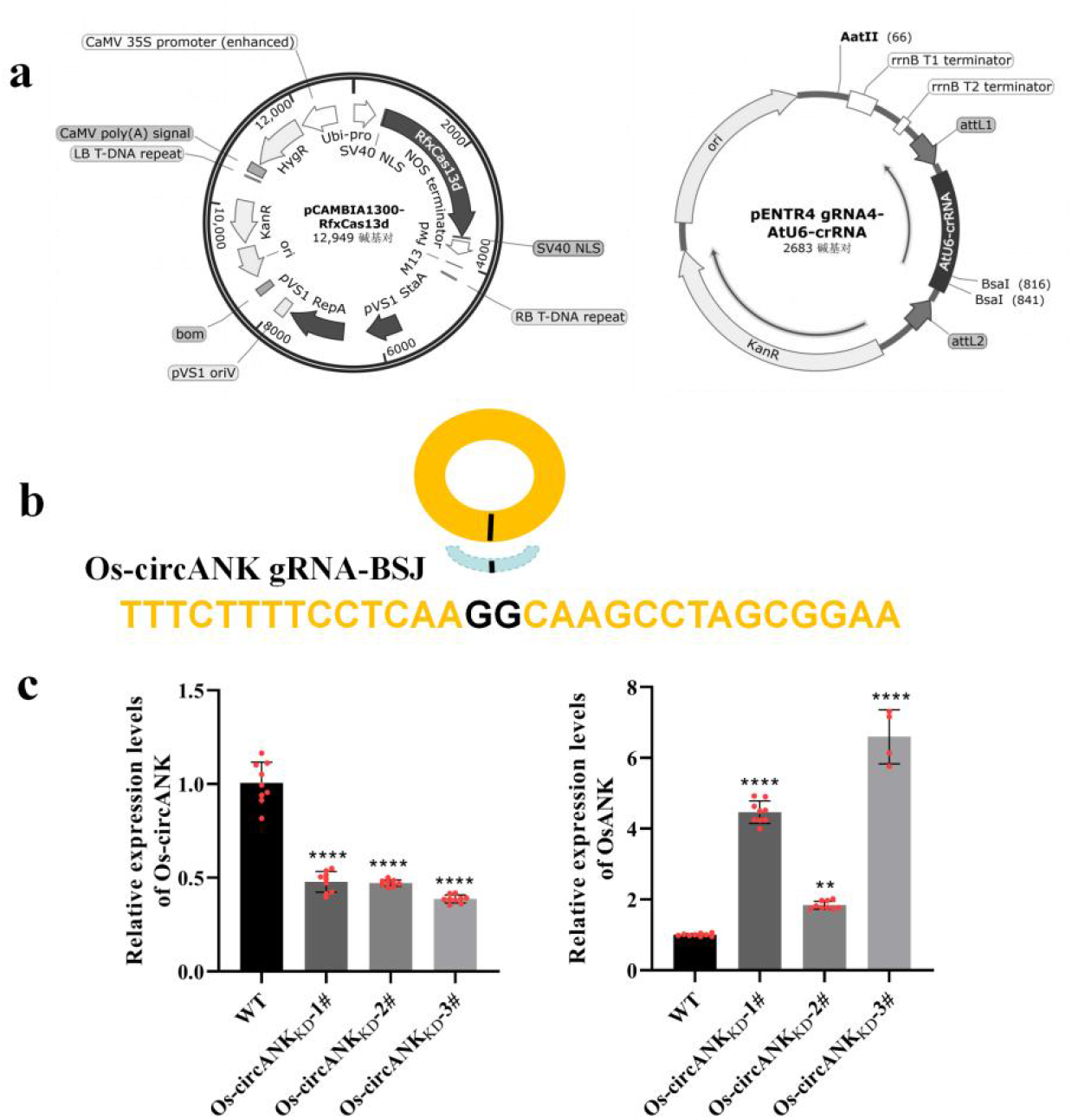
Specific knockdown (KD) of plant circular RNA. a. The schematic of CRISPR-Cas13d vector system. b. The sequence composition of the Os-circANK-specific guide RNA (gRNA) targeting sequences, which specifically encompass the backsplice junction (BSJ) region, is described as follows. c. The expression levels of the Os-circANK and its parental gene (OsANK) in Os-circANK_KD_ lines. *OsUbi* served as a reference gene, mean±SD, t-test, \*\**p* < 0.01, \*\*\*\**p* < 0.0001.

### Os-circANK negatively regulated resistance to bacterial blight in rice

To elucidate the function of Os-circANK against *Xoo*, we generated rice transgenic lines that overexpress Os-circANK (Os-circANK_OE_). qRT-PCR analysis verified the elevated expression of Os-circANK in these transgenic plants (Fig. 3a). To explore the potential interaction of Os-circANK in response to *Xoo* infection, we conducted a disease assay on Os-circANK_OE_ and Os-circANK_KD_ transgenic plants. As a result, Os-circANK_OE_ plants showed increased disease symptoms compared to WT plants, post *Xoo* inoculation, whereas the opposite outcome was demonstrated for the Os-circANK_KD_ plants (Fig. 3b). Leaf-clipping inoculation results demonstrated that Os-circANK_OE_ plants exhibited an average lesion length 2 cm longer than WT plants (Fig. 3b). Accordingly when we examined the bacterial biomass via qPCR, the bacterial biomass in Os-circANK_OE_ plants was significantly higher than that in the WT plants, indicating a faster bacterial growth rate in Os-circANK_OE_ plants (Fig. 3b). Conversely, Os-circANK_KD_ plants showed the opposite results. To further examine whether Os-circANK participates in the regulation of pathogenesis-related (PR) proteins, we evaluated the expression of three PR genes, PR1a^23^, PR1b^24^, and PR10^25^, in the Os-circANK transgenic plants using qRT-PCR. The expression of these three PR genes was upregulated in the Os-circANK_KD_ lines and downregulated in the Os-circANK_OE_ lines compared to the control plants (Fig. 3c). All together, the above results indicate the negative role of Os-circANK in regulating disease resistance in rice.

**Fig.3.**
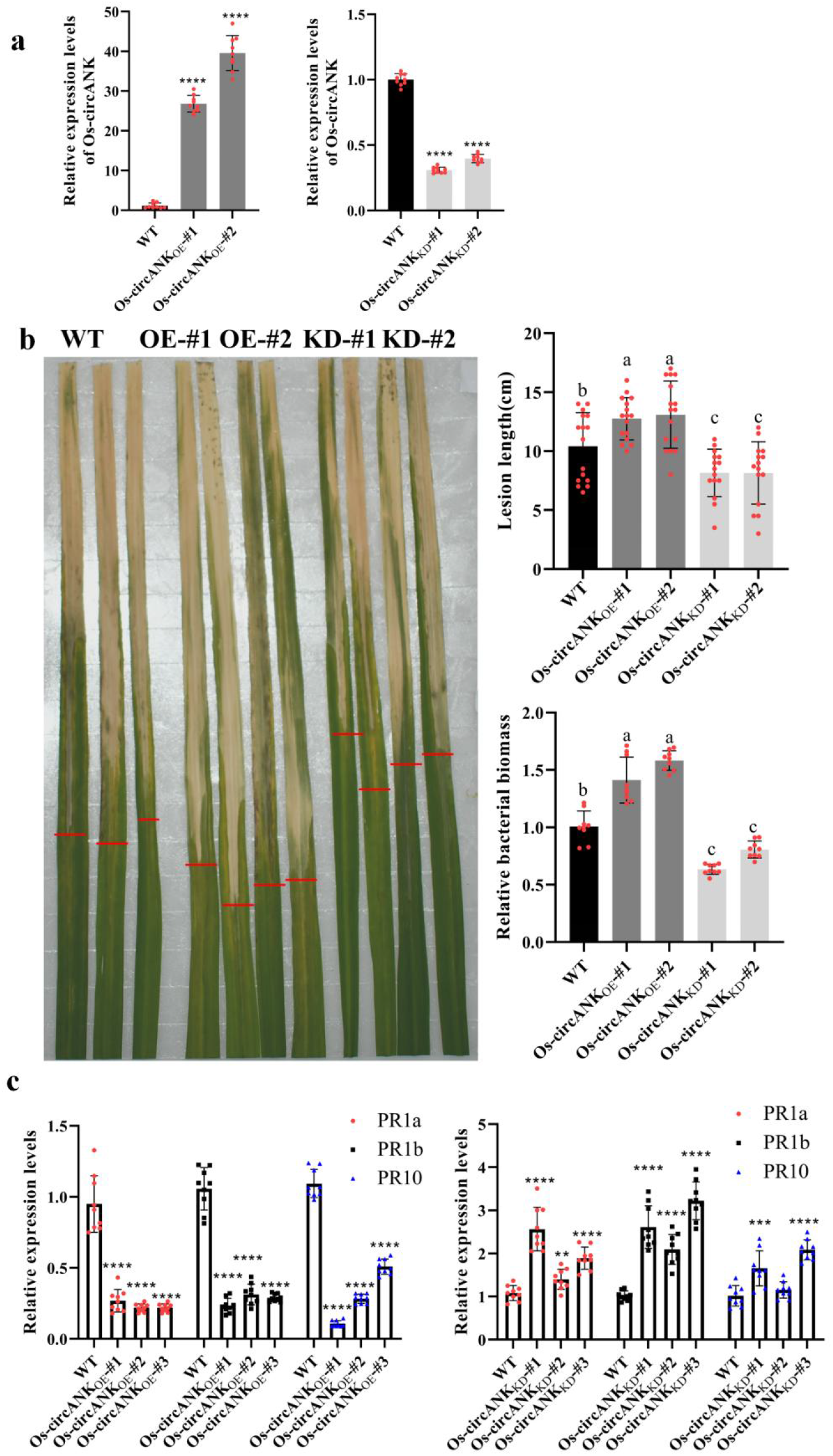
Os-circANK negatively regulated resistance to bacterial blight in rice. a. The expression levels of the Os-circANK in Os-circANK_OE_ and Os-circANK_KD_ lines. b. Disease phenotypes, lesion length and relative bacterial biomass of PXO99^A^-inoculated Os-circANK_OE_, Os-circANK_KD_ and WT plants during the heading stage. c. Relative expression of PR genes (*PR1a*, *PR1b* and *PR10*) in Os-circANK_OE_, Os-circANK_KD_ and WT plants. Values are means±SD of three independent experiments; asterisks represent a significant difference compared to the control group (t-test, \*\**p* < 0.01, \*\*\**p* < 0.001, \*\*\*\**p* < 0.0001). Different letters above the bars indicate significant differences at *p* < 0.01. All of the experiments were repeated three times with similar results.

### Osa-miR398b was identifed as a sponge target of Os-circANK

To determine the coding potential of Os-circANK, a predicted ORF overlapping the BSJ was fused with GUS and introduced into *Nicotiana benthamiana*. None of the fusion constructs exhibited coding activity (Supplementary Fig. S1).

Accumulating evidence substantiates the role of circRNAs as proficient miRNA sponges, adeptly mitigating the inhibitory influence exerted by the latters on their target genes. Simultaneously, cytosolic abundance of Os-circANK suggests its potential role as a competing endogenous RNA (ceRNA). We predicted a common miRNA, Osa-miR398b, through four algorithms (Fig. 4a). The direct interaction between Os-circANK and Osa-miR398b was examined by a luciferase reporter gene assay. Luciferase activity was inhibited by the overexpression of Osa-miR398b in Os-circANK reporter vector containing wild-type Osa-miR398b miRNA recognition element. However, the luciferase expression was not affected in the case of Os-circANK mutated vector (Os-circANK^MUT^) (Fig. 4b). Concurrently, the dual-luciferase assay quantitatively validated these findings (Fig. 4c).

**Fig.4.**
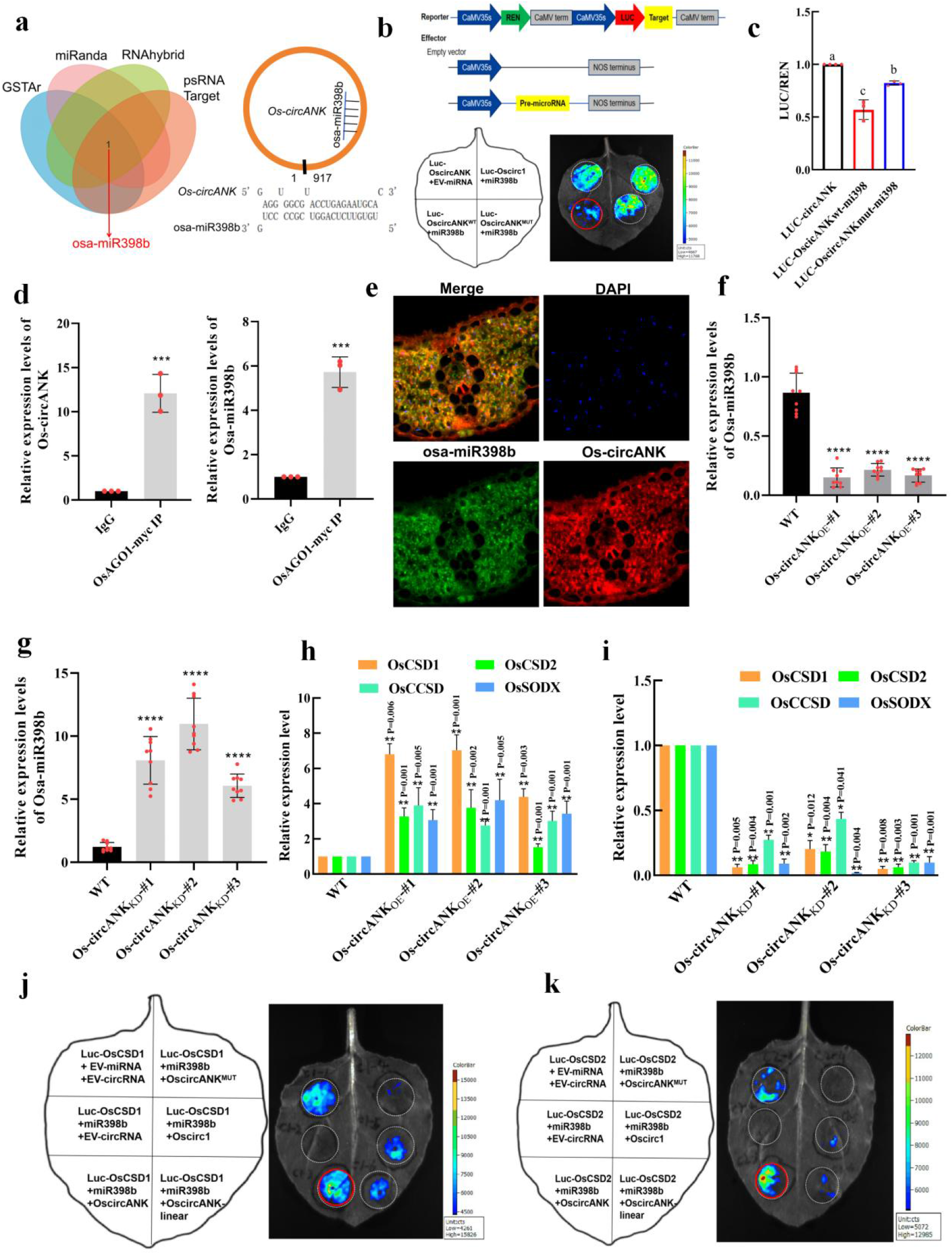
Osa-miR398b is a sponge target of Os-circANK. a. Left panel, a Venn diagram showing Osa-miR398b predicted as a putative sponge target of Os-circANK in four databases. Right panel, predicted Osa-miR398b binding sites within Os-circANK. b,c. To examine the direct interaction between Os-circANK and Osa-miR398b, the luciferase reporter vectors containing the wild-type or mutant sequence of Osa-miR398b binding sites and Os-circ1 were constructed. Os-circ1 was used as a negative control without Osa-miR398b binding sites. *Agrobacterium tumefaciens* EHA105 cells harboring different combinations were used to infiltrate *Nicotiana benthamiana* leaf. Luciferase activities were recorded 3 d after infiltration. Three independent experiments were carried out. Different letters indicate significant differences at *p* < 0.05, as determined by Duncan’s multiple range test. d. The correlation between Osa-miR398b and Os-circANK in rice protoplasts was tested by using RIP assay, anti-myc and anti-IgG antibody were used to immunoprecipitate cellular lysates. e. Determination of the co-localization of Os-circANK and Osa-miR398b by FISH using confocal microscopy. Os-circANK was stained in red, Osa-miR398b was stained in green and nuclei were stained with DAPI. f,g. RT-qPCR analysis of Osa-miR398b in WT, Os-circANK-overexpressing and Os-circANK-knockdown rice leaves. *OsU6* served as a reference gene, mean±SD, t-test, \*\*\*\**p* < 0.0001. h,i. RT-qPCR analysis of *OsCSD1*, *OsCSD2*, *OsCCSD* and *OsSODX* in WT, Os-circANK-overexpressing and Os-circANK-knockdown rice leaves. *OsUbi* served as a reference gene, mean±SD, n = 3, t-test, \**p* < 0.05, \*\**p* < 0.01, \*\*\**p* < 0.001. j,k. The luciferase reporter experiment was implemented to confirm the interaction between Os-circANK, Osa-miR398b and target OsCSD1/OsCSD2. Os-circANK^MUT^ represented mutant Osa-miR398b binding sequence of Os-circANK. Os-circANK-linear represents the forming exons of the Os-circANK, driven only by the 35S promoter. All of the experiments were repeated three times with similar results.

Argonaute (AGO) proteins represent a highly conserved class across eukaryotes, forming complexes with small RNAs (sRNAs) to mediate RNA silencing^26^. In *Arabidopsis thaliana*, AGO1 predominantly associates with miRNAs, exerting post-transcriptional inhibition by cleaving target mRNAs or impeding translation processes within the cytoplasm^26^. Rice possesses four homologous genes designated as OsAGO1a/b/c/d. Previous studies have shown that various defective phenotypes emerge when OsAGO1 is mutated, accompanied by a large accumulation of miRNA target genes^27^. Analysis of sRNA sequences associated with OsAGO1a/b/c proteins showed that OsAGO1 mainly binds to miRNAs with 5’ U base, and most of the miRNAs were distributed in the three OsAGO1s, such as Osa-miR398b^27^.

To validate the interaction between Osa-miR398b and Os-circANK, we constructed an OsAGO1a expression vector with a myc tag and employed transient transformation into rice protoplasts along with ribonucleoprotein immunoprecipitation (RIP) experiments. The results showed that Osa-miR398b and Os-circANK were enriched in the RIP materials compared to the control IgG (Fig. 4d).

Additionally, FISH experiments revealed co-expression of Osa-miR398b and Os-circANK in the cytoplasm (Fig. 4e). In order to examine the transcription levels of Osa-miR398b and its target genes, we performed experiments in Os-circANK overexpressed, knocked down, and control Nipponbare plants. We observed a significant reduction of Osa-miR398b levels in the overexpressed Os-circANK plants and a significant increase of Osa-miR398b levels in the knocked down Os-circANK plants (Fig.4f,g). Conversely, the transcription levels of the Osa-miR398b target genes *OsCSD1, OsCSD2, OsSODX* and *OsCCSD* were increased in the overexpressed Os-circANK plants and decreased in the knocked down Os-circANK plants , compared to the control plants (Fig. 4h,i). Overall, the findings indicate that the cleavage of OsCSD1 and OsCSD2 by Osa-miR398b may be impeded by the presence of the Os-circANK. To validate this interaction, subsequent luciferase reporter gene assays were conducted. The luciferase activity of OsCSD1 and OsCSD2 reporter genes was significantly repressed by Osa-miR398b, with restoration observed only upon addition of Os-circANK and not Os-circ1, linear Os-circANK, or Osa-miR398b binding site-mutated Os-circANK^MUT^ (Fig. 4j,k). The above results indicate that Os-circANK can directly bind to Osa-miR398b and prevent its cleavage activity.

To ascertain the generalizability of miRNA interaction with circRNAs, ath-miR398b and its putative binding circRNA, ath_circ_021952, from *Arabidopsis thaliana* were cloned. After comparing the sequences of Os-circANK and ath_circ_021952, it was found that except for the conserved sequence at the binding site of miR398b, the rest of the sequences are not conserved (Supplementary Fig. S2a). The interaction between ath-miR398b and ath_circ_021952 was validated through a luciferase reporter gene assays (Supplementary Fig. S2b) and observed that the presence of ath_circ_021952 inhibited the cleavage of AtCSD1 and AtCSD2 by ath-miR398b (Supplementary Fig. S2c,d).

### Osa-miR398b and its target genes regulate resistance to bacterial blight in rice

Real-time PCR analysis revealed a notable surge in Osa-miR398b transcription levels post *Xoo* infection in rice, whereas its target genes exhibited a contrasting pattern (Fig. 5a), indicating that Osa-miR398b and its target genes were involved in the interaction between rice and *Xoo*. To investigate the involvement of Osa-miR398b to rice resistance with bacterial diseases, we inoculated the Osa-miR398b overexpressing plants and target mimicry of miR398 (MIM398) line plants at the 12-week-old stage with *Xoo* strain PXO99^A^ using the leaf-clipping method. The disease symptoms were examined at 14 dpi. The average lesion length in the Osa-miR398b overexpressing plants was 8.0 cm, compared to the WT plants and MIM398 lines at 10.4 cm and 12.9 cm, respectively (Fig. 5b). These results show that overexpression of Osa-miR398b confers rice enhanced resistance to bacterial blight disease. Plants with knocked down Os-circANK exhibited phenotypes resembling those observed in plants overexpressing Osa-miR398b.

**Fig.5.**
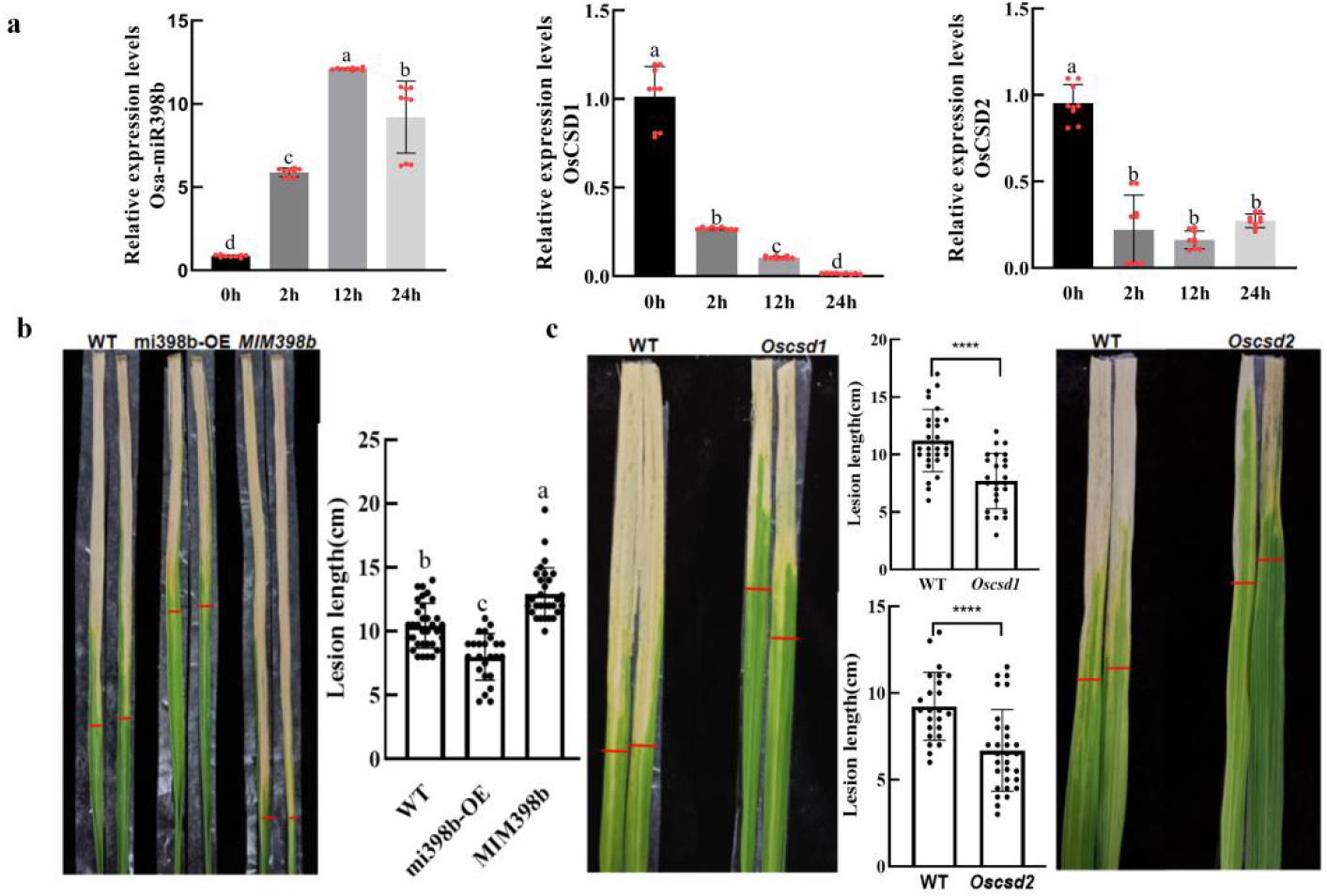
Osa-miR398b and target *OsCSD1*/*OsCSD2* regulated the resistance to bacterial blight in rice. a. RT-qPCR analysis of Osa-miR398b and target *OsCSD1*/*OsCSD2* of PXO99^A^-inoculated Nipponbare plants. b. Disease phenotypes and lesion length on leaves of PXO99^A^-inoculated Osa-miR398b-overexpressing, target mimicry of miR398 (MIM398) and WT plants during the heading stage. c. Disease phenotypes and lesion length on leaves of PXO99^A^-inoculated the indicated mutant lines (Cu/Zn-superoxidase dismutase (csd)1, csd2).

Because a miRNA functions through its target genes, we assessed how Osa-miR398b’s targets, OsCSD1/OsCSD2, affect rice resistance to leaf blight disease. The mean lesion lengths in the mutant strains Oscsd1 and Oscsd2 were 7.7 cm and 6.7 cm, respectively, in contrast to the WT, which exhibited lesion lengths of 11.0 cm and

9.2 cm, respectively (Fig. 5c).

### Os-circANK regulate ROS concentration upon *Xoo* infection by Osa-miR398b and its target genes

Rapid ROS production is a crucial indicator of activated plant defense systems. We assayed H_2_O_2_ and O·^2-^ concentrations in the Os-circANK overexpressing and knockdowned lines under mock and *Xoo* treatments. Consistent with disease phenotypes, H_2_O_2_ concentration rose significantly in Os-circANK_KD_ upon PXO99^A^ infection compared to the WT plants, but decreased and significantly lower in Os-circANK_OE_ (Fig. 6a). While there was no significant difference in O·^2-^ levels between Os-circANK_KD_ and WT, both were higher than in Os-circANK_OE_ lines (Fig. 6b). We subsequently assayed the mRNA levels of three NADPH oxidase genes: RbohB (Nagano et al., 2016), RbohD, and RbohF (Jang et al., 2012). Compared to the WT plants, the transcription levels of three NADPH oxidase genes were significantly upregulated in the Os-circANK_KD_ plants, while Os-circANK_OE_ plants exhibited an opposite trend (Fig. 6c).

**Fig.6.**
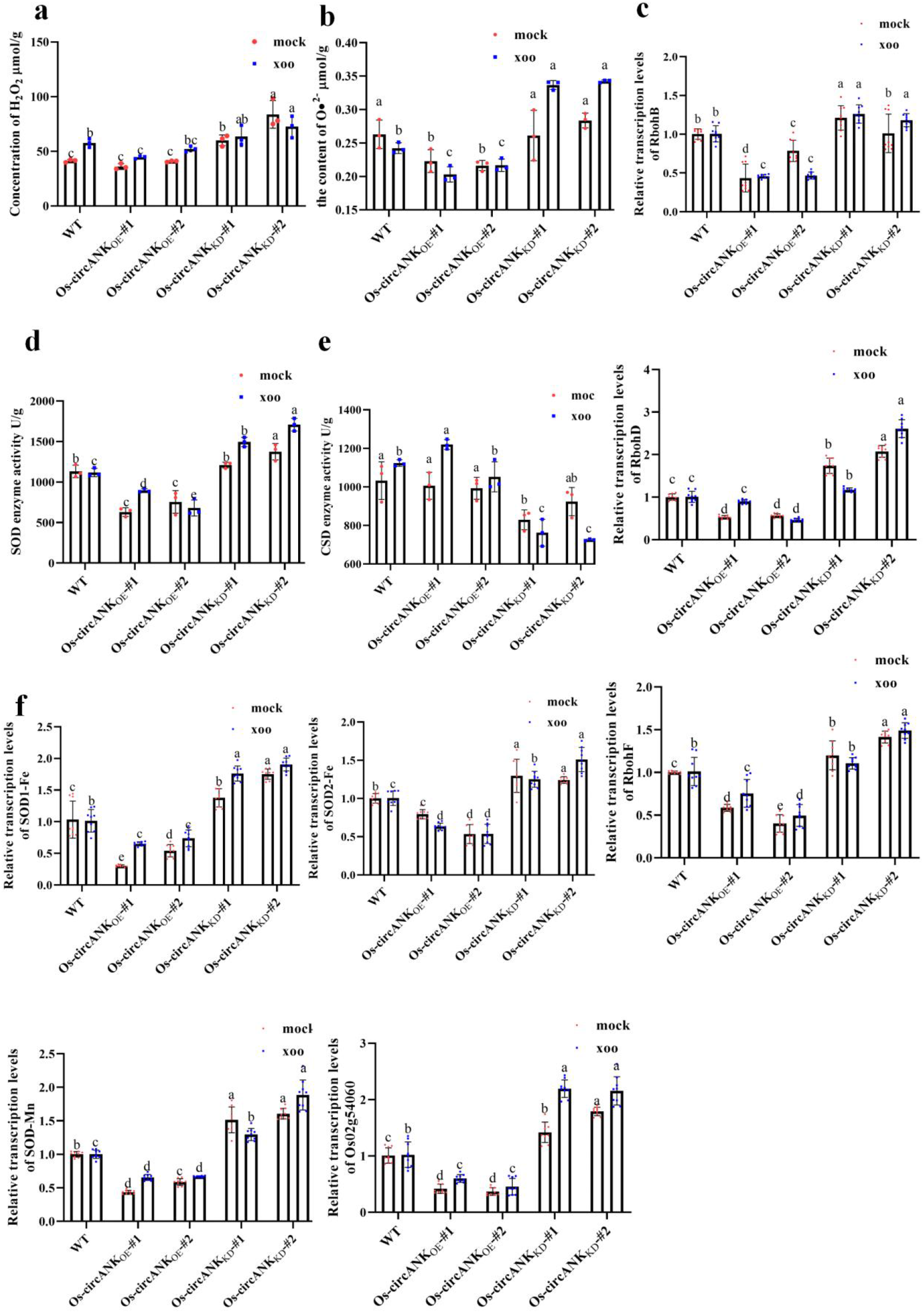
Os-circANK affected reactive oxygen species (ROS) concentration, Cu/Zn-Superoxidase Dismutase (CSD) and total Superoxidase Dismutase (SOD) enzyme activity upon *Xoo* infection. a, b Quantification of hydrogen peroxide (H_2_O_2_) (a) and superoxide radicals (O^•^_2_^−^) (b) concentrations in leaves of the Os-circANK_OE_, Os-circANK_KD_ and WT plants with PXO99^A^/mock treatment, respectively. Values are means of three replications. Error bars indicate SD. Different letters above the bars indicate significant differences at *p* < 0.01. c. RT-qPCR data showing the transcription levels pattern of the indicated NADPH oxidase genes (Respiratory burst oxidase homologs(Rboh)B, RbohD and RbohF) in the indicated lines upon PXO99^A^/mock treatment. d. The total SOD enzyme activity in transgenic lines and WT plants with PXO99^A^/mock treatment. e. The enzyme activity of CSD in indicated lines with PXO99^A^/mock treatment. f. RT-qPCR data showing the transcription levels pattern of the SOD family genes in the indicated lines upon PXO99^A^/mock treatment. Relative mRNA amounts were normalized to that in WT mock samples. Different letters above the bars indicate significant differences at *p* < 0.01. All of the experiments were repeated two times with similar results.

Subsequently, we assayed the total SOD and CSD enzyme activity in all tested lines. Our data showed that upon *Xoo* infection, the total SOD enzyme activity was significantly higher in the Os-circANK_KD_ lines compared to both the WT and Os-circANK_OE_ lines (Fig. 6d). On the contrary, the CSD enzyme activity was significantly reduced in the Os-circANK_KD_ lines (Fig. 6e). Subsequently, the mRNA levels of other SOD family genes (*SOD1-Fe*, *SOD2-Fe*, *SOD-Mn*, and a SOD family chaperone, *Os02g54060*) were examined. Consistently with the results of SOD levels, the transcription levels of these genes was significantly upregulated in the Os-circANK_KD_ lines (Fig. 6f). Overall, these results indicate that upon *Xoo* infection, downregulated Os-circANK enhances the Osa-miR398b-mediated degradation of OsCSDs, which in turn triggers the upregulation of other SOD family members resulting in higher total SOD enzyme activity, leading to higher ROS concentration and enhanced blight disease-resistance.

## Discussion

CircRNA is a covalently closed single-stranded RNA generated by reverse splicing of exons from precursor mRNA (Jeck and Sharpless, 2014). Past studies on plant circRNAs primarily concentrated on comprehensive genome identification, with limited functional characterizations of circRNAs (Lu et al., 2015; Xiang et al., 2018; Zhou et al., 2018; Wang et al., 2019b; Wang et al., 2022). With the exception of the back-splicing junction site, the majority of circRNAs exhibit complete sequence overlap with their homologous linear RNA (Liu and Chen, 2022). This presents challenges in the computational detection, experimental validation, and functional assessment of circRNAs. In plants, the CRISPR-Cas9 system has been employed to suppress circRNA biogenesis either by directly knocking out the exons forming the circRNAs or by disrupting the reverse complementary sequences in the flanking regions (Zhou et al., 2021). However, this approach carries the risk of off-target effects, potentially resulting in the deletion of the parental gene (Zhou et al., 2021). Li *et al*. discovered that the knockdown efficiency of circRNAs achieved through RfxCas13d-BSJ-gRNA averages 65%, surpassing the 35% efficiency observed with shRNA (Li et al., 2022). This suggests that CRISPR-RfxCas13d holds significant potential for widespread application in the investigation of circRNA function. Therefore, we designed and constructed a CRISPR-RfxCas13d vector specifically tailored for the targeted knockdown of plant circRNAs. By utilizing gRNA to target the back-splicing junction of Os-circANK, this system effectively distinguishes the circRNA from the linear mRNA. It demonstrates the capability for the specific depletion of Os-circANK without influencing the expression of its homologous linear RNA (Fig. 2c). The development of this method provides a valuable tool for functional studies of plant circRNAs.

We identified a downregulated circRNA, Os-circANK, following *Xoo* infection in rice (Fig. 1). Through overexpression and knockdown experiments, we validated the negative regulatory role of Os-circANK in rice resistance to *Xoo* (Fig. 3). These findings broaden our understanding of the functional roles of circRNAs in the plant-pathogens interactions. Apart from our study, only one research has indicated that a circRNA can modulate rice resistance to blast fungus through overexpression techniques (Fan et al., 2020). Rice lines overexpressing circR5g05160 exhibited smaller lesion sizes and reduced fungal biomass upon infection with blast fungus. This suggests that circR5g05160 enhances resistance to blast fungus in rice (Fan et al., 2020). Furthermore, there are studies that indicate the role of circRNAs in abiotic stress responses. The overexpression of Vv-circPTCD1 negatively impacted growth under heat, salt, and drought stresses in Arabidopsis (Ren et al., 2023). These findings indicate that circRNAs contribute to the response of plants to various stresses.

Furthermore, several studies in animals have demonstrated that circRNAs can sponge miRNAs, thereby upregulating the expression of their target genes (Hansen et al., 2013). Similarly, extensive predictions and construction of circRNA-miRNA-mRNA expression networks using bioinformatics tools suggest the existence of ceRNA mechanisms in plant circRNAs (Ye et al., 2019; Zhang et al., 2020). The most direct experimental evidence involves altering the expression of circRNAs and examining whether the expression of miRNAs and their target mRNAs is subsequently affected. In this study, overexpression and knockdown of Os-circANK demonstrated its ability to bind to Osa-miR398b, resulting in reduced expression of Osa-miR398b and consequently increased expression of its target genes, OsCSD1/OsCSD2 (Fig. 4f,j,h,i). Further validation of the Os-circANK and Osa-miR398b binding was confirmed through Luciferase reporter assays and RIP (Fig. 4c,d). The establishment of the regulatory relationship among Os-circANK-Osa-miR398b-OsCSD1/OsCSD2 provides novel evidence supporting the role of plant circRNA as a ceRNA.

The rapid generation of ROS, such as O^•^_2_^−^ and H_2_O_2_, serves as a pivotal indicator for activating the plant defense system (Gill and Tuteja, 2010). Rbohs play a pivotal role by catalyzing the transfer of electrons from NADPH to oxygen, resulting in the generation of O^•^_2_^−^ (Kaur et al., 2014). SOD isoenzymes are categorized into three types based on their metal ion affinity in plants: Cu/Zn-Superoxide Dismutase (CSD), manganese SOD (SOD-Mn), and iron SOD (SOD-Fe) (Mittler, 2002). SODs play a crucial role as primary defense components against various stresses by modulating the H_2_O_2_ concentration within plants (Mittler, 2002). MiR398 is a conserved microRNA family that inhibits the expression of members belonging to the SOD family across multiple plant species (Guan et al., 2013). MiR398 and CSDs play crucial roles in regulating plant disease resistance against pathogens. In rice, the overexpression of miR398b is associated with increased basal defenses against *M. oryzae* (Li et al., 2019). This enhanced resistance is linked to reduced mRNA levels of *CSD1*, and *CSD2* (Guan et al., 2013). In our study, we evaluated H_2_O_2_, O·^2-^ levels and total SOD enzyme activity in transgenic lines with Os-circANK_KD_ or Os-circANK_OE_ following *Xoo* inoculation. Consistently, our findings demonstrated that Os-circANK_KD_ lines exhibited elevated ROS levels and enhanced total SOD enzyme activity(Fig. 6), leading to enhanced resistance to blight disease. In conclusion, these findings lay the groundwork for exploring the role of circRNAs in the interaction between plants and pathogens.

In summary, upon *Xoo* infection, downregulation of Os-circANK occurs, leading to an upregulation of miR398b. Consequently, miR398b inhibits the expression of *CSD1* and *CSD2*, resulting in an overall elevation of SOD enzyme activity. This cascade of events culminates with the accumulation of H_2_O_2_, thereby strengthening the resistance of rice to *Xoo* through an enhanced oxidative defense mechanism (Fig. 7).

**Fig.7.**
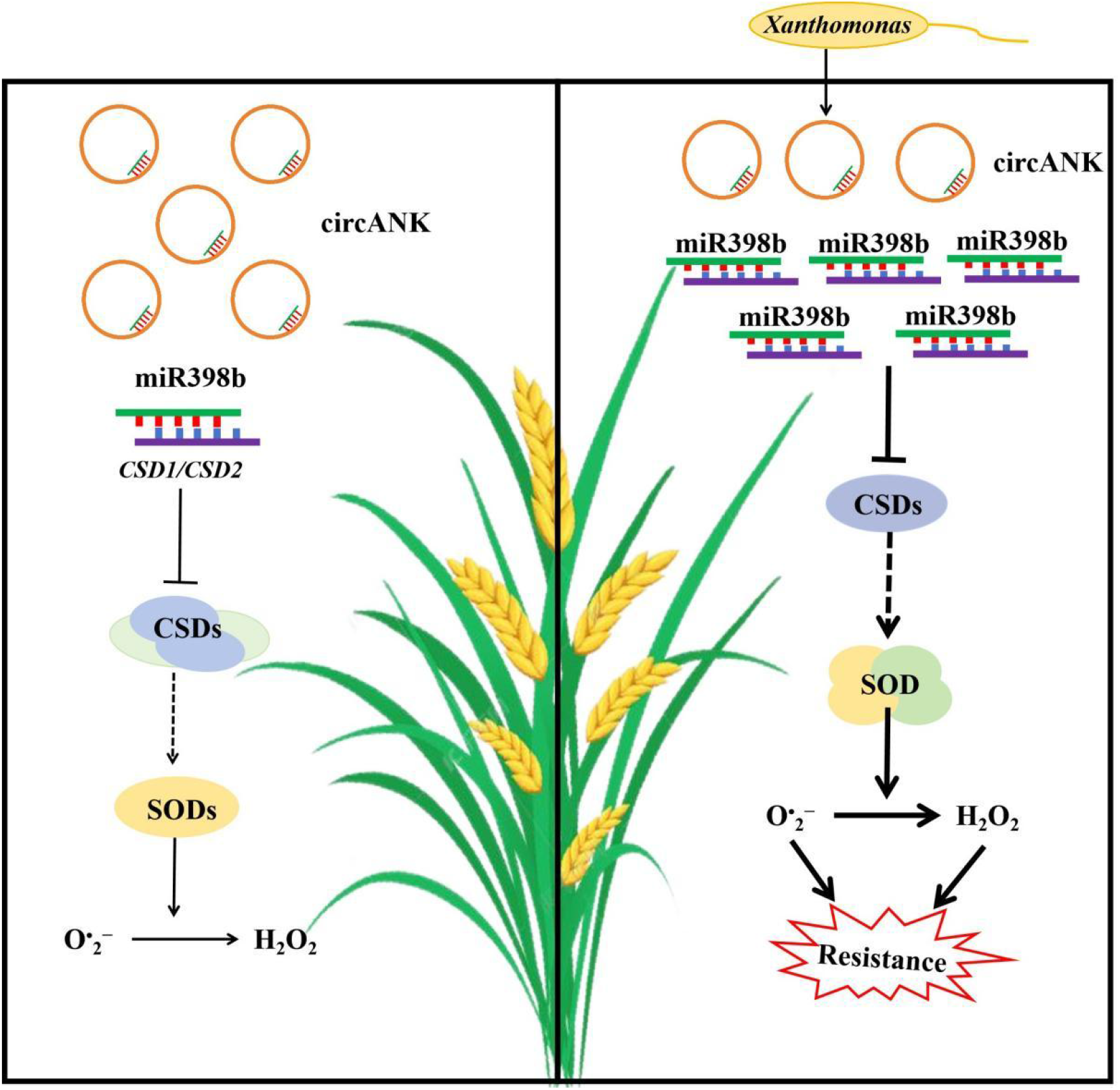
A model of Os-circANK role in plant immunity against *Xoo*. In the absence of pathogen infection, Os-circANK modulates OsCSD1/CSD2 expression by competitively binding Osa-miR398b, maintaining basal CSD and total SOD enzyme activity in plant cells. A compensatory mechanism between CSD and other SODs regulates hydrogen peroxide concentrations for normal cellular metabolism. Upon *Xoo* infection, decreased Os-circANK expression increases Osa-miR398b levels, leading to CSD1/CSD2 repression and reduced CSD activity. This triggers compensatory upregulation of other SODs, enhancing total SOD enzyme activity and promoting hydrogen peroxide synthesis, thus increasing resistance. Arrows in the diagram represent positive regulation, while blunt-ended bars indicate inhibition. The dotted lines indicate the unidentified regulation that exists between CSDs and SODs.

## Methods

### Plant materials

The rice (*Oryza sativa*) japonica accessions Nipponbare (NPB) was used for transgenic analysis. For *Xoo*-resistance and resistant response assays, the wild-type (WT) control and the Os-circANK_OE_ (overexpression of Os-circANK) and Os-circANK_KD_ (downregulation of Os-circANK) transgenic lines were grown in a greenhouse at 28±2℃ and 70% relative humidity under 12 h : 12 h, light : dark cycles. The laboratory of Professor Wenming Wang at Sichuan Agricultural University supplied the seeds of the transgenic lines, including the Osa-miR398b overexpressing line, target mimicry of miR398 (MIM398) line and the mutant lines with suppressed *Oscsd1*/*Oscsd2* expression. All seeds underwent a germination period of 5 days in distilled water and were then transplanted into soil, where they were cultivated in a greenhouse.

### Construction of CRISPR/Cas13d system for knocking down plant circRNA

The gene sequence of RfxCas13d was subjected to optimization according to the codon bias of plants and subsequently synthesized. The optimized sequence was cloned into a linearized pUC57 vector, designated as pUC57:RfxCas13d. Subsequently, pUC57:RfxCas13d and a CRISPR/spCas9 vector were subjected to double digestion with *Bam*H I and *Nco* I restriction endonucleases. The resulting 3003 bp RfxCas13d gene fragment and the CRISPR/spCas9 vector backbone of approximately 12.6 kb were independently recovered. The 3003 bp RfxCas13d fragment was then ligated to the CRISPR/spCas9 vector backbone using T4 DNA ligase, yielding the recombinant plasmid named pCAMBIA1300-RfxCas13d.

The pENTR4:gRNA4 vector was enzymatically linearized using *Hind* III, and subsequently, the linearized vector backbone was isolated. The AtU6-crRNA gene fragment, featuring two *Bsa* I cleavage sites, was synthesized through artificial means and then ligated into the *Hind* III cleavage site of the pENTR4:gRNA4 vector. This molecular manipulation resulted in the generation of the pENTR4:gRNA4-AtU6-crRNA vector.

The targeting sequence of guide RNAs (gRNAs) was designed based on the BSJ site of circRNAs, and primers corresponding to the RfxCas13d-BSJ-gRNA target sequence were synthesized. Subsequently, these primers were annealed and ligated into the *Bsa* I linearized pENTR4:gRNA4-AtU6-crRNA vector. The resulting construct, named pENTR4:gRNA4-AtU6-BSJ-crRNA vector, served as a template for primer design, incorporating *Hind* III and *Sbf* I cleavage sites to amplify the AtU6-BSJ-crRNA fragments. The pCAMBIA1300-RfxCas13d recombinant vector underwent double digestion with *Hind* III and *Sbf* I. Ultimately, the pCAMBIA1300-RfxCas13d-BSJ-crRNA vector was obtained through homologous recombination.

Agrobacterium strain EHA105 was selected for rice genetic transformation. Hygromycin B was used for screening the genotype of transgenic plants by means of hygromycin resistance analysis.

### RT-qPCR analysis

For mRNA and circRNA quantification, RNA was reverse-transcribed using six-base random primers. For miR398b quantification, stem-loop primers were designed. All the primers used are shown in Supporting Information Supplementary Table S1.

### RNase R digestion, nucleoplasmic separation, and RNA-FISH

Total RNA extracted from ‘Nipponbare’ leaves was incubated at 37°C for 30 min with 2 U RNase R (Geneseed) and then purified using an RNA clean kit (DP412; Tiangen, China). Nucleoplasmic separation experiments and RNA-FISH in rice cells were performed as previously described (Gao et al., 2019; Nielsen et al., 2022). Probes with Cy3 modification for Os-circANK and FAM-labeled probes specific to miR398b were synthesized by Service Biotech (Wuhan, China). Nuclei were stained with DAPI and photographed by confocal microscopy.

### Prediction of miRNAs for Os-circANK interactions

Four bioinformatic prediction algorithms (GSTAr.pl, https://github.com/MikeAxtell/GSTAr; miRanda, http://www.microrna.org; RNAhybrid, http://bibiserv.techfak.uni-bielefeld.de/rnahybrid/; and psRNATarget, https://www.zhaolab.org/psRNATarget/) were used to search for Os-circANK-interacting miRNAs.

### Dual-luciferase reporter assay

For analysis of the circRNA-microRNA interaction, WT and mutated binding site sequences of Os-circANK were cloned into pGreenII 0800-miRNA vectors, respectively. The pre-miR398b sequences were amplified from rice genomic DNA and inserted into a pCambia1301 vector.

For analysis of circRNA-microRNA-mRNA interaction, an overexpression vector for Os-circANK was generated by amplifying the Os-circANK sequence, including its flanking intron regions (500 bp upstream and downstream), from rice genomic DNA.

Subsequently, the amplified sequences were inserted between reverse complementary segments into the pHB vector (Gao et al., 2019). For the linear fragment expression vector, the linear fragment derived from the same sequence with Os-circANK was cloned into the pHB vector. The microRNA target sequences with a size of 200-300 nt were amplified from rice cDNA and then inserted into the pGreenII 0800-miRNA vector.

The procedures for introducing the assembled vectors into *Agrobacterium tumefaciens* and the subsequent suspension, infiltration, and cultivation of tobacco plants were conducted following established methodologies outlined in a previous study (Gao et al., 2023).

Luciferase signals were assayed using an IVIS Lumina II system, following manufacturer’s protocol. Relative luciferase intensities were determined using a Dual-luciferase reporter assay Kit (Promega), according to standard operating procedures.

### Rice Protoplast Preparation, Transfection and Ribonucleoprotein immunoprecipitation (RIP)

Rice protoplast isolation was conducted following the protocol described by Zhang et al. (Zhang et al., 2014) with minor modifications. Briefly, 10-day-old rice shoots were sliced into ∼0.5 mm strips and incubated in an enzyme solution consisting of 1% cellulase R-10 and 0.4% macerozyme R-10 for 4-6 hours in the dark under gentle agitation (80 rpm). Subsequently, the pellets were rinsed with wash buffer and centrifuged at 100 g for 10 minutes to collect the rice protoplasts.

The *OsAGO1a* sequences with a myc tag were amplified from rice cDNA, and inserted into a pCambia1301 vector. Finally, rice protoplasts (100 µl) were transfected with 10 µg of *35S*_pro_: OsAGO1a-myc plasmids.

Anti-IgG, anti-myc antibodies were incubated with magnetic beads at 4 ℃ for 2h to overnight according to the instructions of the PureBinding®RNA Immunoprecipitation Kit (Geneseed). Subsequently, the antibody-bead complex underwent overnight incubation at 4 °C with cell lysates derived from rice protoplasts. The resulting bound RNA was eluted, reverse transcribed into cDNA, and subsequently quantified through qRT-PCR.

### Disease assay

Freshly activated *Xoo* strain PXO99^A^ was inoculated into liquid NB medium (5 g/L peptone, 10 g/L sucrose, 1 g/L Yeast Extract, and 3 g/L beef extract) and shaken overnight for cultivation. Eight weeks-old rice plants were infected with *Xoo* through the scissor-dip method using bacterial suspensions at OD_600_ = 0.8. The disease incidence on leaves was assessed at 14 days post-inoculation (dpi). The qPCR to analyse relative bacterial biomass, which was represented by the ratio of the *Xoo* gene (Sun et al., 2024) to the Ubiquitin gene of rice.

## Acknowledgements

We thank Professor Wenming Wang and Yan Li (Sichuan Agricultural University) for providing the seeds of the transgenic lines, including the Osa-miR398b overexpressing line, target mimicry of miR398 (MIM398) line and the mutant lines with suppressed Oscsd1/Oscsd2 expression. We thank Professor Fangfang Li and Xueping Zhou (The Institute of Plant Protection, Chinese Academy of Agricultural Sciences) for providing the SpCas9–1301 plasmid.

## Author contributions

B.Z., Z.X. and G.C conceived the study and designed the experiments. S.W. and R.L. performed data analyses. X.L., X.F., Z.Z., S.L., W.L., M.O., and P.W. performed the experiments. P.S. and V.M. revised the manuscript. X.L. wrote the paper, and all authors commented on the paper.

## Competing interests

The authors declare no competing interests.

All remaining data are in the main paper or the supplementary materials. Source data are provided with this paper.

---

**Supplementary Fig. S1.**
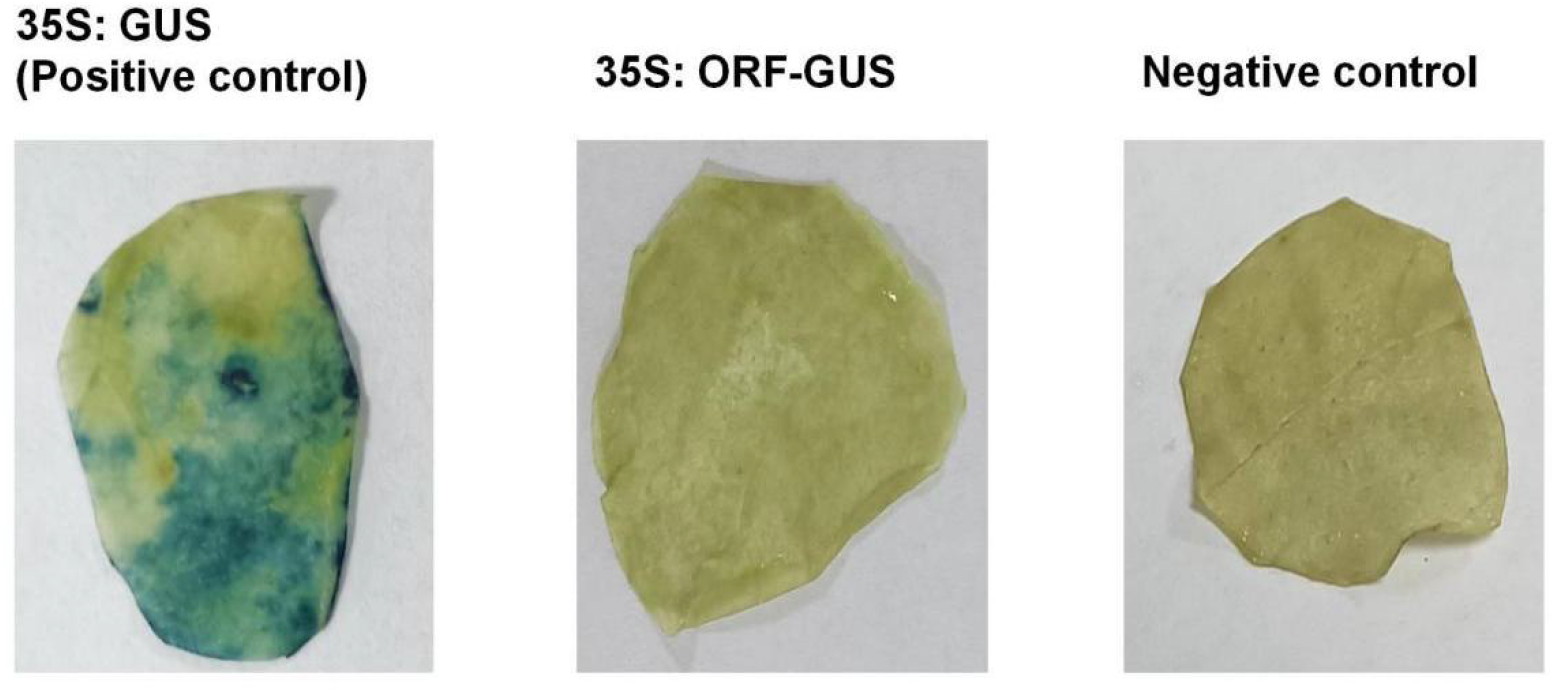
Os-circANK coding potential analysis. ORF Finder analysis indicates that Os-circANK has a potential ORF spanning reverse splicing sites starting from ATG. The 35S::ORF-GUS vector fused the upstream sequence and ORF without the stop codon with the downstream reporter GUS gene. The empty vector GUS-1301 was used as positive control, and transformed to *N. benthamiana* leaves. Untransformed leaves as negative control. β-Glucuronidase (GUS) staining was used to assess the potential coding capacity.

**Supplementary Fig. S2.**
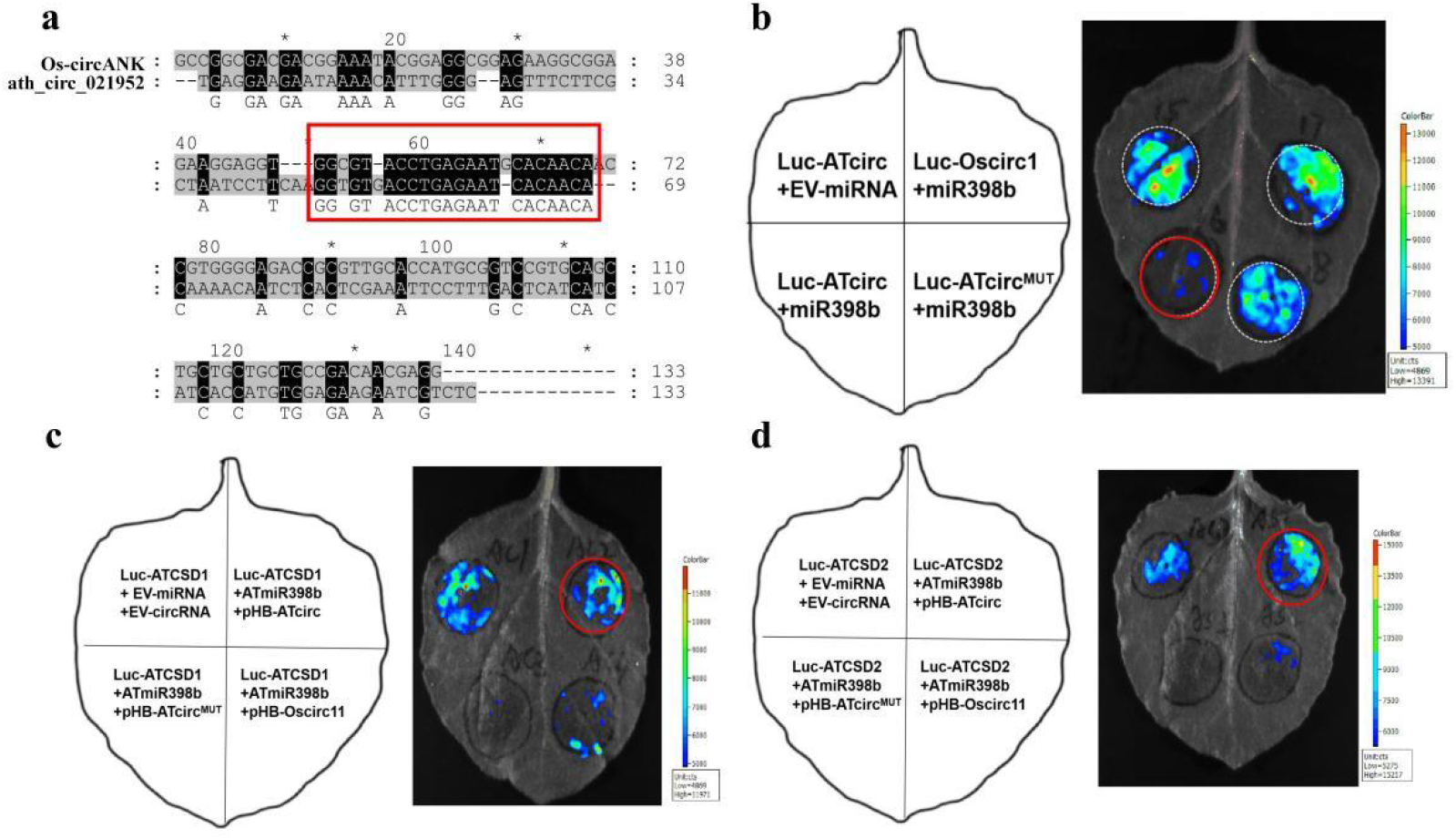
Experimental verification of ath_circ_021952 competitive binding to ath-miR398b. a, b. The interaction between ath-miR398b (AtmiR398) and ath_circ_021952 (Atcirc) was validated through a luciferase reporter gene assays. c,d. The luciferase reporter experiment was implemented to confirm the interaction between ath_circ_021952 (Atcirc), ath-miR398b (AtmiR398) and target AtCSD1/AtCSD2. Atcirc^MUT^ represented mutant AtmiR398b binding sequence of Atcirc. Oscirc11 as negative control.

## References

Busungu, C., Taura, S., Sakagami, J.I., and Ichitani, K. (2016). Identification and linkage analysis of a new rice bacterial blight resistance gene from XM14, a mutant line from IR24. Breed Sci 66, 636–645.

Chen, L., Wang, C., Sun, H., Wang, J., Liang, Y., Wang, Y., and Wong, G. (2021). The bioinformatics toolbox for circRNA discovery and analysis. Brief Bioinform 22, 1706–1728.

Chen, L., Zhang, P., Fan, Y., Lu, Q., Li, Q., Yan, J., Muehlbauer, G.J., Schnable, P.S., Dai, M., and Li, L. (2018). Circular RNAs mediated by transposons are associated with transcriptomic and phenotypic variation in maize. New Phytol 217, 1292–1306.

Fan, J., Quan, W., Li, G.B., Hu, X.H., Wang, Q., Wang, H., Li, X.P., Luo, X., Feng, Q., Hu, Z.J., Feng, H., Pu, M., Zhao, J.Q., Huang, Y.Y., Li, Y., Zhang, Y., and Wang, W.M. (2020). circRNAs Are Involved in the Rice-Magnaporthe oryzae Interaction. Plant Physiol 182, 272–286.

Gao, Z., Sun, B., Fan, Z., Su, Y., Zheng, C., Chen, W., Yao, Y., Ma, C., and Du, Y. (2023). Vv-circSIZ1 mediated by pre-mRNA processing machinery contributes to salt tolerance. New Phytol 240, 644–662.

Gao, Z., Li, J., Luo, M., Li, H., Chen, Q.J., Wang, L., Song, S.R., Zhao, L.P., Xu, W.P., Zhang, C.X., Wang, S.P., and Ma, C. (2019). Characterization and Cloning of Grape Circular RNAs Identified the Cold Resistance-Related Vv-circATS1. Plant Physiol 180, 966–985.

Gill, S.S., and Tuteja, N. (2010). Reactive oxygen species and antioxidant machinery in abiotic stress tolerance in crop plants. Plant Physiol Biochem 48, 909–930.

Guan, Q., Lu, X., Zeng, H., Zhang, Y., and Zhu, J. (2013). Heat stress induction of miR398 triggers a regulatory loop that is critical for thermotolerance in Arabidopsis. Plant J 74, 840–851.

Guo, J.U., Agarwal, V., Guo, H., and Bartel, D.P. (2014). Expanded identification and characterization of mammalian circular RNAs. Genome Biol 15, 409.

Han, Y., Li, X.X., Yan, Y., Duan, M.H., and Xu, J.H. (2020). Identification, characterization, and functional prediction of circular RNAs in maize. Mol Genet Genomics 295, 491–503.

Hansen, T.B., Jensen, T.I., Clausen, B.H., Bramsen, J.B., Finsen, B., Damgaard, C.K., and Kjems, J. (2013). Natural RNA circles function as efficient microRNA sponges. Nature 495, 384–388.

Hong, Y.H., Meng, J., Zhang, M., and Luan, Y.S. (2020). Identification of tomato circular RNAs responsive to Phytophthora infestans. Gene 746.

Huang, J., Zhou, W.L., Zhang, X.M., and Li, Y. (2023). Roles of long non-coding RNAs in plant immunity. Plos Pathog 19.

Jeck, W.R., and Sharpless, N.E. (2014). Detecting and characterizing circular RNAs. Nat Biotechnol 32, 453–461.

Kaur, G., Sharma, A., Guruprasad, K., and Pati, P.K. (2014). Versatile roles of plant NADPH oxidases and emerging concepts. Biotechnol Adv 32, 551–563.

Kristensen, L.S., Andersen, M.S., Stagsted, L.V.W., Ebbesen, K.K., Hansen, T.B., and Kjems, J. (2019). The biogenesis, biology and characterization of circular RNAs. Nat Rev Genet 20, 675–691.

Li, S.Q., Wu, H., and Chen, L.L. (2022). Screening circular RNAs with functional potential using the RfxCas13d/BSJ-gRNA system. Nat Protoc 17, 2085–2107.

Li, Y., Cao, X.L., Zhu, Y., Yang, X.M., Zhang, K.N., Xiao, Z.Y., Wang, H., Zhao, J.H., Zhang, L.L., Li, G.B., Zheng, Y.P., Fan, J., Wang, J., Chen, X.Q., Wu, X.J., Zhao, J.Q., Dong, O.X., Chen, X.W., Chern, M., and Wang, W.M. (2019). Osa-miR398b boosts H(2) O(2) production and rice blast disease-resistance via multiple superoxide dismutases. New Phytol 222, 1507–1522.

Liu, C.X., and Chen, L.L. (2022). Circular RNAs: Characterization, cellular roles, and applications. Cell 185, 2016–2034.

Liu, S., Wu, L., Qi, H., and Xu, M. (2019). LncRNA/circRNA–miRNA–mRNA networks regulate the development of root and shoot meristems of Populus. Industrial Crops and Products 133, 333–347.

Lu, T., Cui, L., Zhou, Y., Zhu, C., Fan, D., Gong, H., Zhao, Q., Zhou, C., Zhao, Y., Lu, D., Luo, J., Wang, Y., Tian, Q., Feng, Q., Huang, T., and Han, B. (2015). Transcriptome-wide investigation of circular RNAs in rice. RNA 21, 2076–2087.

Mittler, R. (2002). Oxidative stress, antioxidants and stress tolerance. Trends Plant Sci 7, 405–410.

Nielsen, A.F., Bindereif, A., Bozzoni, I., Hanan, M., Hansen, T.B., Irimia, M., Kadener, S., Kristensen, L.S., Legnini, I., Morlando, M., Jarlstad Olesen, M.T., Pasterkamp, R.J., Preibisch, S., Rajewsky, N., Suenkel, C., and Kjems, J. (2022). Best practice standards for circular RNA research. Nat Methods 19, 1208–1220.

Nino-Liu, D.O., Ronald, P.C., and Bogdanove, A.J. (2006). Xanthomonas oryzae pathovars: model pathogens of a model crop. Mol Plant Pathol 7, 303–324.

Ren, Y., Li, J., Liu, J., Zhang, Z., Song, Y., Fan, D., Liu, M., Zhang, L., Xu, Y., Guo, D., He, J., Song, S., Gao, Z., and Ma, C. (2023). Functional Differences of Grapevine Circular RNA Vv-circPTCD1 in Arabidopsis and Grapevine Callus under Abiotic Stress. Plants (Basel) 12.

Sun, Q.P., Xiao, Y.X., Song, L., Yang, L., Wang, Y., Yang, W., Yang, Q., Xie, K.B., Yuan, M., and Li, G.T. (2024). Mutation of confers broad-spectrum disease resistance in rice. Molecular Plant Pathology 25.

Wang, P., Wang, S., Wu, Y., Nie, W., Yiming, A., Huang, J., Ahmad, I., Zhu, B., and Chen, G. (2022). Identification and Characterization of Rice Circular RNAs Responding to Xanthomonas oryzae pv. oryzae Invasion. Phytopathology 112, 492–500.

Wang, P.L., Bao, Y., Yee, M.C., Barrett, S.P., Hogan, G.J., Olsen, M.N., Dinneny, J.R., Brown, P.O., and Salzman, J. (2014). Circular RNA is expressed across the eukaryotic tree of life. Plos One 9, e90859.

Wang, Y., Xiong, Z., Li, Q., Sun, Y., Jin, J., Chen, H., Zou, Y., Huang, X., and Ding, Y. (2019a). Circular RNA profiling of the rice photo-thermosensitive genic male sterile line Wuxiang S reveals circRNA involved in the fertility transition. Bmc Plant Biol 19, 340.

Wang, Y., Gao, Y., Zhang, H., Wang, H., Liu, X., Xu, X., Zhang, Z., Kohnen, M.V., Hu, K., Wang, H., Xi, F., Zhao, L., Lin, C., and Gu, L. (2019b). Genome-Wide Profiling of Circular RNAs in the Rapidly Growing Shoots of Moso Bamboo (Phyllostachys edulis). Plant Cell Physiol 60, 1354–1373.

Xiang, L.X., Cai, C.W., Cheng, J.R., Wang, L., Wu, C.F., Shi, Y.Z., Luo, J.Z., He, L., Deng, Y.S., Zhang, X., Yuan, Y.L., and Cai, Y.F. (2018). Identification of circularRNAs and their targets in Gossypium under Verticillium wilt stress based on RNA-seq. Peerj 6.

Ye, J., Wang, L., Li, S., Zhang, Q., Zhang, Q., Tang, W., Wang, K., Song, K., Sablok, G., Sun, X., and Zhao, H. (2019). AtCircDB: a tissue-specific database for Arabidopsis circular RNAs. Brief Bioinform 20, 58–65.

Zhang, J., Hao, Z., Yin, S., and Li, G. (2020). GreenCircRNA: a database for plant circRNAs that act as miRNA decoys. Database (Oxford) 2020.

Zhang, Y.C., Liao, J.Y., Li, Z.Y., Yu, Y., Zhang, J.P., Li, Q.F., Qu, L.H., Shu, W.S., and Chen, Y.Q. (2014). Genome-wide screening and functional analysis identify a large number of long noncoding RNAs involved in the sexual reproduction of rice. Genome Biol 15, 512.

Zhou, J.P., Yuan, M.Z., Zhao, Y.X., Quan, Q., Yu, D., Yang, H., Tang, X., Xin, X.H., Cai, G.Z., Qian, Q., Qi, Y.P., and Zhang, Y. (2021). Efficient deletion of multiple circle RNA loci by CRISPR-Cas9 reveals Os06circ02797 as a putative sponge for OsMIR408 in rice. Plant Biotechnol J 19, 1240–1252.

Zhou, R., Zhu, Y.X., Zhao, J., Fang, Z.W., Wang, S.P., Yin, J.L., Chu, Z.H., and Ma, D.F. (2018). Transcriptome-Wide Identification and Characterization of Potato Circular RNAs in Response to Pectobacterium carotovorum Subspecies brasiliense Infection. Int J Mol Sci 19.

